# ERK activation in CAR T cells is amplified by CD28-mediated increase in CD3ζ phosphorylation

**DOI:** 10.1101/718767

**Authors:** Jennifer A. Rohrs, Elizabeth L. Siegler, Pin Wang, Stacey D. Finley

## Abstract

Chimeric antigen receptors (CARs) are engineered receptors that mediate T cell activation. CARs are comprised of activating and costimulatory intracellular signaling domains derived from endogenous T cells that initiate signaling required for T cell activation, including ERK activation through the MAPK pathway. Understanding the mechanisms by which co-stimulatory domains influence signaling can help guide the design of next-generation CARs. Therefore, we constructed an experimentally-validated computational model of anti-CD19 CARs in T cells bearing the CD3ζ domain alone or in combination with CD28. We used ensemble modeling to explore different mechanisms of CD28 co-stimulation on the ERK response time. Model simulations show that CD28 primarily influences ERK activation by enhancing the phosphorylation kinetics of CD3ζ, predictions that are validated by experimental measurements. Overall, we present a mechanistic mathematical modeling framework that can be used to gain insights into the mechanism of CAR T cell activation and produce new testable hypotheses.

## 1 INTRODUCTION

Chimeric antigen receptor (CAR) engineered T cells have recently been approved for the treatment of CD19+ cell malignancies (Mullard, 2017). These therapies have been extremely successful for CD19+ hematological cancers, but it has been difficult to extend CAR T cell therapies to other types of cancer, specifically solid tumors (Morgan et al., 2010). To better engineer CAR T cells to fight cancer, we need to improve our understanding of how these modified receptors activate T cells.

CARs typically include an extracellular antibody-derived binding domain linked to a transmembrane domain and a number of different intracellular signaling domains (Sadelain, Brentjens, & Rivière, 2013). These signaling domains are derived from endogenous T cells and typically include CD3ζ, a part of the endogenous T cell receptor (TCR), and a costimulatory domain, such as CD28. It is clear that T cells require this secondary signaling through a co-stimulatory receptor, but the mechanisms through which co-stimulatory domains influence T cell activation are not clear (Bretscher, 1999). Additionally, it is not clear how CAR signaling differs from endogenous T cell receptor signaling (Harris et al., 2018).

Computational mechanistic models can be used to test hypotheses about molecular signaling mechanisms. These models have been used in the past to study endogenous T cell activation, providing insights into important activation and feedback mechanisms that help control the sensitivity and specificity of TCR activation, reviewed previously (Rohrs, Wang, & Finley, 2019). These models generally assume that T cell activation is derived directly from the TCR CD3ζ signaling domain, while neglecting the effects of the co-stimulatory domains. Therefore, the immunology field has developed a fairly clear picture of the signaling events downstream of CD3ζ, but there is a lack of understanding of the effects of co-stimulation.

Recently, we have used phospho-proteomic mass spectrometry combined with mechanistic computational modeling to gain more insight into the effects of CAR co-stimulation. We quantified the site-specific phosphorylation kinetics of CARs containing CD3ζ with or without CD28 (referred to as 28z and Z, respectively) (Rohrs et al., 2018). Our experimental data showed that CD3ζ immunoreceptor tyrosine-based activation motifs (ITAMs) are phosphorylated independently, in a random order, and with distinct kinetics. Adding the CD28 co-stimulatory domain increased the rate of CD3ζ phosphorylation by over 3-fold. In addition, by applying the model, we identified that LCK phosphorylates CD3ζ through a mechanism of competitive inhibition. However, our integrative experimental and modeling approach does not explain how the increase in CD3ζ phosphorylation affects downstream signaling. More generally, it is not clear how any of the effects of CD28 influence downstream T cell activation. We are particularly interested in activation of the MAPK signaling pathway, leading to ERK phosphorylation, as this pathway exhibits clear switch-like response in T cells and helps mediate T cell activation and proliferation (Altan-Bonnet & Germain, 2005).

To explain how the CAR intracellular domains influence ERK response time, we constructed a mechanistic computational model of T cell activation by CARs containing the CD3ζ domain alone or in combination with CD28. We first calibrate the model using published experimental data and show that the model is able to reproduce known effects of various intracellular protein perturbations on ERK response time following T cell activation. We then use an ensemble modeling approach to predict the effects of various mechanisms of CD28 co-stimulation (Brännmark, Palmér, Glad, Cedersund, & Strålfors, 2010). Experimental measurements of ERK response in CAR-engineered T cells validate the model hypothesis that CD28 activates ERK primarily through modifications of CD3ζ phosphorylation kinetics. The model also generates additional hypotheses that can be used to guide new experiments. Overall, this modeling study enriches our understanding of CAR T cell co-stimulatory activation.

## 2 METHODS

### Construction of a mechanistic computational model of CAR T cell activation

We constructed a model of CAR T cell activation based on our previous modeling work, as well as other models and experimentally measured kinetic data and parameters in the literature. Overall, the model presented in this work includes signaling initiated by antigen binding to the CAR and culminates in phosphorylation of ERK. The full model (**Figure 1**) includes models of kinetic proofreading (Altan-Bonnet & Germain, 2005; Coombs & Goldstein, 2005; McKeithan, 1995) and kinetic segregation (Barua, Faeder, & Haugh, 2007; Davis & van der Merwe, 2006), CD45 phosphatase activity (Arulraj & Barik, 2018), LCK autoregulation (Rohrs, Wang, & Finley, 2016), CAR phosphorylation (Rohrs et al., 2018), LAT signalosome formation, Ras activation (Das et al., 2009), MAPK pathway activation (Birtwistle et al., 2012), and SHP1 negative feedback (Altan-Bonnet & Germain, 2005). The steps used to unite these elements into a single model are described below.

**Figure 1.**
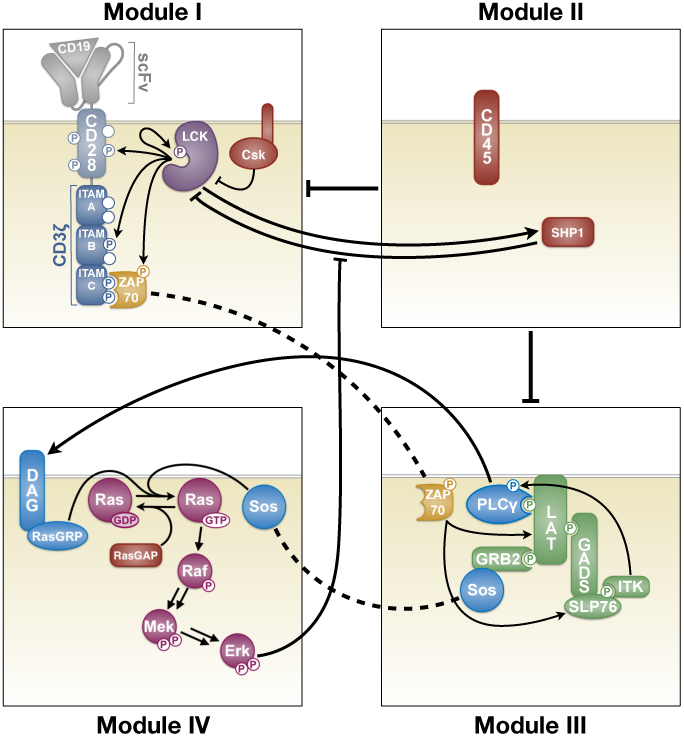
Schematic of signaling model from CAR antigen binding through ERK activation, incorporating models from literature. Arrow and bars indicate activating and inhibitory interactions, respectively. Dashed lines denote the same species in multiple Modules. **Module I**: LCK regulation, autophosphorylation, and phosphorylation of the CAR intracellular signaling domains and ZAP-70. **Module II:** CD45 and SHP1 phosphatase activity. CD45 is constitutively active in resting T cells, but it is excluded from the signaling area upon CAR binding to its ligand. SHP1 is recruited to the area by singly phosphorylated CD3ζ ITAMs and is activated by LCK. **Module III:** The LAT signalosome forms when ZAP-70 binds to doubly phosphorylated ITAMs and becomes phosphorylated by LCK. It can then phosphorylate sites on LAT and SLP76. Phosphorylated sites on LAT can bind proteins Grb2, GADS, and PLCg. Grb2 can bind to SOS, while GADS binds to SLP76. SLP76 recruits Tec family kinases, like ITK, which can then phosphorylate and activate PLCg. **Module IV:** PLCg and SOS can activate Ras-GDP to Ras-GTP. Ras-GTP can be inactivated by RasGAP. Once activated, RAS-GTP can activate the MAPK pathway, which leads to ERK activation. Doubly phosphorylated ERK can phosphorylate LCK at a protection site, which prevents LCK from associating with SHP1, resulting in a positive feedback loop.

#### LCK autoregulation

We have previously developed a mass action based model of LCK autoregulation (Rohrs, Wang, et al., 2016). However, due to the size of the complete CAR signaling model, we simplified the model of LCK autoregulation to reduce the computational complexity. To do this, we altered the interactions between various phosphorylated forms of LCK from mass action kinetics to Michaelis-Menten kinetics and added in the significant protein-protein binding reactions identified by the original LCK auto-regulation model. This greatly reduced the number of ordinary differential equations, as all pairs of different phosphorylated LCK species no longer need to form dimers before the autophosphorylation reactions can be catalyzed. The parameters from this newly reduced minimal LCK model were refit to the same data used to parameterize the original model (Hui & Vale, 2014). In the fitting process, it was determined that pairs of Michaelis-Menten constants and catalytic rates as well as pairs of association and dissociation rates were not independently identifiable. Therefore, Michaelis-Menten constants were held at a value of 1000 molecules/µm^2^, which is in line with previously fit values from similar systems (Rohrs, Zheng, Graham, Wang, & Finley, 2018) and within the range of protein concentrations in the model. Association rates were held constant at a value of 0.1 µm^2^molecule^-1^min^-1^ (Northrup & Erickson, 1992; Schlosshauer & Baker, 2004). These assumptions for the values of Michalis-Menten constants and association rates were used throughout the model when explicit literature values were unavailable. Fitting was done using particle swarm optimization (PSO), described below.

#### CAR phosphorylation and kinetic proofreading

Before stimulation, LCK autophosphorylation is allowed to reach steady state in the presence of the phosphatase CD45. Antigen is then added to the model and allowed to bind to the CAR. The dissociation constant for binding of the antigen to the CAR was hand-tuned to agree with the *in vitro* 28ζ CAR T cell ERK activation experimental data measured in this work, described below.

Antigen-bound CAR can be phosphorylated by active LCK, as we have quantified and modeled previously. The parameters governing these interactions were adapted directly from our previous work (Rohrs et al., 2018). We assume that only LCK phosphorylated on the activating site, Y394, is catalytically active toward the CAR and other downstream proteins in the T cell activation pathway (Philipsen et al., 2017). As our previously published model of CAR tyrosine site phosphorylation only accounts for the catalytic activity of active LCK monophosphorylated at Y394, we assumed that doubly phosphorylated LCK, phosphorylated at the activating site Y394 and the inhibitory site Y505, has a catalytic activity 50x slower than active monophosphorylated LCK. Kinetic proofreading occurs when antigens with lower affinities unbind from the CAR. Free tyrosine sites on this antigen-unbound CAR can then be dephosphorylated by the phosphatase CD45.

#### Kinetic segregation

Binding of an antigen to a TCR or CAR protein leads to a narrow region between the T cell and target cell that excludes the large extracellular domain of CD45 (Leupin, Zaru, Laroche, Müller, & Valitutti, 2000; Mukherjee, Mace, Carisey, Ahmed, & Orange, 2017; Watanabe, Kuramitsu, Posey, & June, 2018). This is modeled by a transition of CD45 from an accessible form to an inaccessible form based on the relative amount of antigen-bound and -unbound CAR. It has been shown that CD45 is excluded from the immunological synapse and that this synapse formation occurs between 5-30 minutes after T cell engagement with a target cell (Huppa & Davis, 2003); therefore, we assume that the CD45 transport has a half-life of 30 minutes, and its exclusion rate is proportional to the amount of antigen-bound CD3ζ.

#### ZAP activation

ZAP-70 is a kinase that can bind to doubly phosphorylated ITAMs on CD3ζ. ZAP binding protects these ITAM sites from dephosphorylation. The parameters governing the binding of ZAP to the ITAMs were adapted from experimental measurements in the literature (Bu, Shaw, & Chanti, 1995; Katz, Novotná, Blount, & Lillemeier, 2017). Additionally, the kinase activity of CAR-bound ZAP can be activated through phosphorylation by LCK, and we assume that this occurs with a slightly slower catalytic rate than LCK phosphorylation of CD3ζ ITAMs. Active ZAP can then unbind from the CD3ζ ITAMs and subsequently phosphorylate downstream proteins in the LAT signalosome. As we do not explicitly model the spatial heterogeneity and movement of ZAP from the ITAM to the LAT signalosome, we assume that ZAP phosphorylates its substrates 10-fold slower than the catalytic rate of LCK phosphorylation of ZAP.

#### LAT signalosome

In the LAT signalosome, we include only proteins that directly relate to the activation of the ERK and the MAPK pathway (Braiman, Barda-Saad, Sommers, & Samelson, 2006; Brownlie & Zamoyska, 2013; Nag, Monine, Faeder, & Goldstein, 2009). All of the dissociation constants governing these interactions were taken from measurements in the literature (Houtman et al., 2004).

#### CD28 activation

CD28 contains four tyrosine sites that are able to be phosphorylated by LCK (Rohrs et al., 2018). These sites bind a variety of downstream signaling proteins, similar to the LAT signalosome. Dissociation constants for these interactions were taken from the literature (Higo et al., 2014; Tian et al., 2015).

#### Ras activation

The mechanism of Ras-GTP activation was adapted from a model by Das *et al*. (Das et al., 2009). The activators of Ras-GTP from Ras-GDP are SOS and RAS-GRP. In our model, SOS is able to bind directly to Grb2 in the LAT signalosome and on CD28. Ras-GRP, is activated by DAG, produced by cleavage of PIP2 by PLCg. PLCg can be activated by Tec family kinases, which are recruited by SLP76 binding in the LAT signalosome and on CD28. Little quantitative information is known about the parameters governing Tec family kinase activity; however, it has been shown that the activity of these kinases is directly related to SLP76 binding (Bogin, Ainey, Beach, & Yablonski, 2007). Therefore, we assumed that the rate of PLCg activation is proportional to the amount of SLP76 bound to the CAR signaling region on LAT or CD28.

#### MAPK pathway

The MAPK pathway and its parameters were directly adapted from the Birtwistle *et al*. model, using the zero feedback case (F=1) (Birtwistle et al., 2012). This pathway includes the three-layer phosphorylation cascade involving RAF, MEK, and ERK.

#### SHP1 negative feedback

The last mechanism we included in the model was negative feedback through the phosphatase SHP1, which can be turned off by positive feedback from activated ERK, first modeled by Altan-Bonnet and Germain (Altan-Bonnet & Germain, 2005). The catalytic rate parameters governing SHP1 activation and its activity were assumed to be rapid, but slightly slower than LCK phosphorylation of CD3ζ. The association rate of SHP1 was tuned to agree with the 28ζ *in vitro* CAR T cell ERK activation experimental data described below.

### Initial conditions

Initial concentrations of proteins in the model were adapted from values calculated in the literature or previous models (Birtwistle et al., 2012; Das et al., 2009; Hui et al., 2017; Hui & Vale, 2014; Lipniacki, Hat, Faeder, & Hlavacek, 2008). In the model, all interactions are assumed to take place in the region on or near the T cell membrane; therefore, for ease of comparison and calculation, all species concentrations were converted to units of molecules/µm^2^. To covert from parameter values and concentrations in units per volume to units per surface area, we assume an average T cell radius of 6 µm. Volume was calculated assuming a spherical shape, and surface area was calculated assuming a sphere with a roughness factor of 1.8, as described previously (Hui & Vale, 2014).

### Parameter fitting using particle swarm optimization (PSO)

The minimal model of LCK autoregulation was fit to data using a particle swarm optimization (PSO) algorithm (Iadevaia, Lu, Morales, Mills, & Ram, 2010). We allowed the catalytic rates to vary on a log scale from 10^-1^-10^4^ min^-1^. The objective function was calculated to minimize the sum of the squared errors between the model outputs and the experimental data. The PSO algorithm was run 100 times, and the median value of each parameter was selected for use in the model.

We also fit the catalytic rates of CD45 dephosphorylation to data from Hui *et al.* 2017 (Hui et al., 2017). A similar method using this data was used previously to explore the effects of PD1 on T cell signaling (Arulraj & Barik, 2018). To do this, we simulated our model using only the species included in the experiments performed by Hui and coworkers. The initial conditions for these species were set according to the equivalent molecules/µm^2^ value listed in the Hui *et al*. supplemental information. All other species’ initial concentrations in our model were set to 0. We then recorded the model outputs of the normalized phosphorylation of various species at 30 minutes for a range of different CD45 concentrations, mimicking the experiments performed by Hui *et al*. We fit the model 100 times using PSO, keeping all Michaelis-Menten constants (K_M_) equal to 1000 molecules/µm^2^, a value on the same order of magnitude as the protein concentrations and the average K_M_ values from the LCK minimal model.

This fitting procedure provided a set of parameter values that enabled the model to match the available experimental data (**Figure 2**). All model parameters and their values are listed in Supplemental File S1. Initial concentrations are provided in Supplemental File S2, and the final fitted model is given in Supplemental File S3.

**Figure 2.**
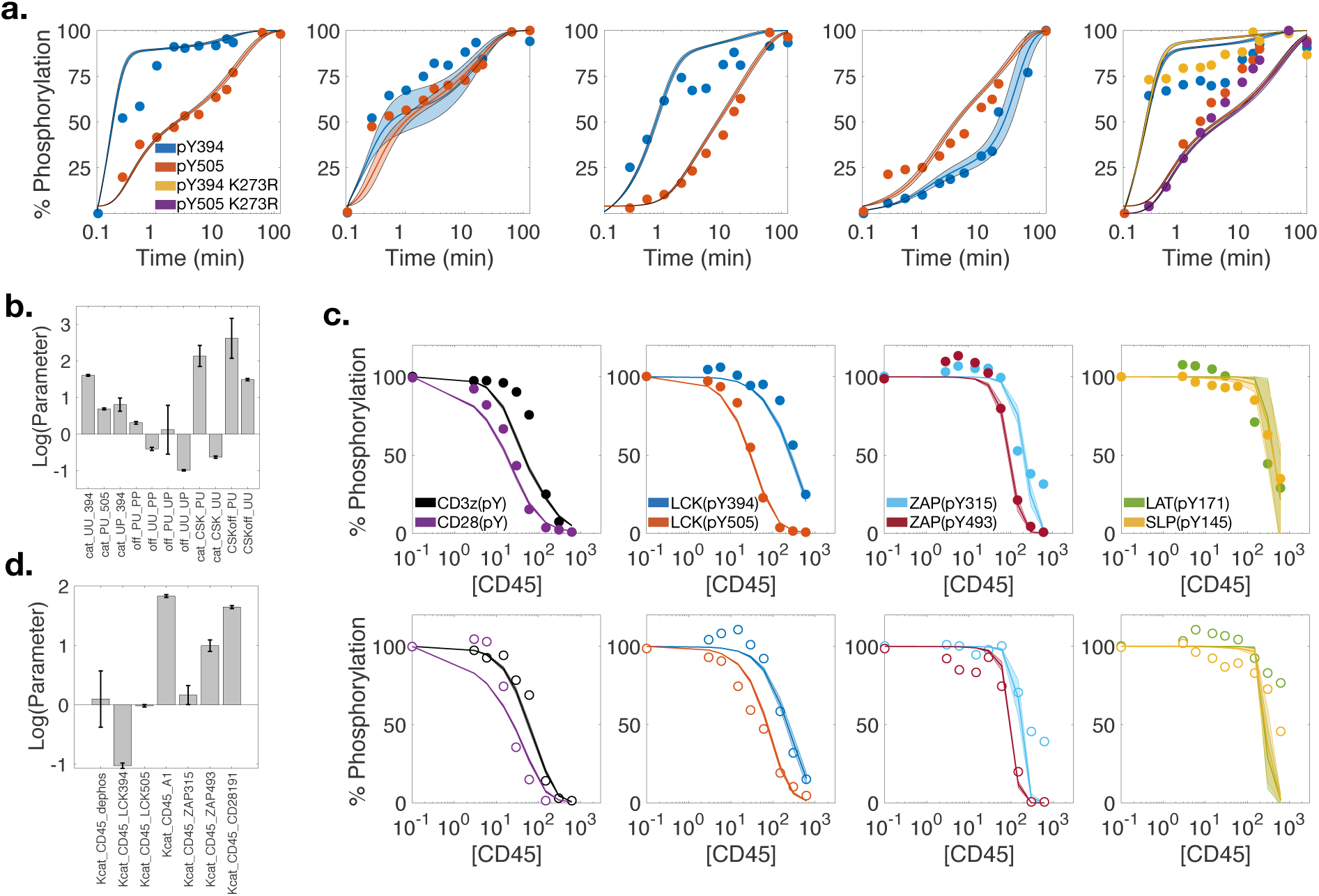
Model parameters were fit to experimental data. a) A reduced version of our model of LCK regulation (Rohrs, Wang, et al., 2016) was refit to data (dots) from Hui and Vale (Hui & Vale, 2014) for five different experimental conditions using Michaelis-Menten kinetics for all LCK-LCK catalytic interactions and mass action kinetics for all CSK-LCK interactions and LCK-LCK binding interactions. The median value from 100 fitted parameter sets (solid lines) is shown, with the shaded region indicating the standard deviation of the fits. b) The median value and standard deviation of the fitted LCK and CSK parameter values from 100 optimized sets (on log scale). LCK catalytic parameters are denoted as cat_XX_#, where XX indicates the phosphorylation state of Y394 and Y505 on the substrate LCK and # indicates the tyrosine site substrate being phosphorylated. Catalytic parameters for CSK phosphorylation of LCK Y505 are denoted as cat_CSK_XX, where XX indicates the phosphorylation state of the LCK substrate. Dissociation rates are denoted as off_XX_YY, where XX and YY indicate the phosphorylation state of the LCK binding partners at Y394 and Y505, respectively, or CSKoff_XX where the binding partners are CSK and LCK phosphorylated at Y394 or Y505 as indicated by XX, respectively. All fitted parameters are in units of min^-1^. c) *Top* row, CD45 catalytic rate parameters were fit to data from Hui et al. in the absence of CSK (Hui et al., 2017) (dots). The median value of 100 optimized parameter sets (solid lines) is shown, with the shaded region indicating the standard deviation. *Bottom row*, as validation, the model was simulated with 145 molecules/µm^2^ CSK and compared to data from Hui et al. not used in parameter fitting (open circles). The median (lines) and standard deviation (shaded region) of the 100 optimized parameter sets is shown. d) The median value and standard deviation of the CD45 catalytic parameter values from 100 optimized sets (shown on a log scale). CD45 catalytic rates are denoted as Kcat_CD45_x, where x indicates the substrate tyrosine sites. A1 indicates CD3ζ ITAM tyrosine sites. Dephos is a generic dephosphorylation rate applied to all substrates not specifically fit to their own value. Catalytic rates are in units of min^-1^.

### Sensitivity analysis

The extended Fourier amplitude sensitivity test (eFAST) has been described in detail previously (Marino, Hogue, Ray, & Kirschner, 2008), and we have applied this approach in our previous work (Rohrs, Sulistio, & Finley, 2016; Rohrs, Wang, et al., 2016). Briefly, eFAST is a global variance-based sensitivity analysis that can identify which model parameters have the most significant effect on a given model output. In this method, a set of model parameters are varied at the same time, with different frequencies, and the model output is calculated. The Fourier transform of the model output is then compared to the various frequencies with which the parameters were varied. A model output’s sensitivity to a given parameter of interest is proportional to the normalized Fourier transform peak of the model output at the frequency with which that parameter was varied. This is referred to as the individual sensitivity index (*S*_*i*_). The extent of higher order interactions between parameters can then be estimated by calculating the Fourier transform peaks of frequencies other than those of the individual frequency and harmonics of the parameter of interest, giving the total sensitivity index (*S*_*Ti*_). A greater total index compared to the first-order index indicates that a parameter is more important in combination with other parameters than alone. The effect of a parameter is considered to be statistically significant if its sensitivity index is greater than that of a dummy variable.

We implemented the eFAST method using MATLAB code developed by Kirschner and colleagues (Marino et al., 2008). We analyzed the parameters in seven groups, allowing each parameter to vary 10-fold up and down from its baseline value.

### Cell lines and reagents

Jurkat T cells (ATTC TIB-152) and K562 cells (ATCC CCL-243) were maintained in 5% CO_2_ environment in RPMI (GIBCO) media supplemented with 10% fetal bovine serum, 1% penicillin-streptomycin, and 2 mM L-glutamine. Alexa-488 conjugated antibody against doubly phosphorylated ERK (clone E10) was purchased from Cell Signaling Technology. Anti-HA antibody (AB9110) was purchased from Abcam. Alexa-647 conjugated goat anti-rabbit secondary antibody was purchased from Thermo Scientific. PE conjugated anti-CD19 (clone HIB19) was purchased from Biolegend.

### Stable transductions of CAR-and CD19-expressing cell lines

The construction of a lentiviral plasmid containing an HA-tagged anti-CD19 CAR bearing the CD28 transmembrane domain and CD28 and CD3ζ intracellular domains (28z) was described previously (Siriwon et al., 2018). Briefly, the anti-CD19 sequence was based on a previously reported CD19 CAR (Milone et al., 2009). The codon optimized CD19 single-chain fragment variable (scFv) sequence and human CD8 hinge region (aa 138-184) was synthesized by Integrated DNA Technologies (Coralville, IA). The CD19/CD8 hinge gene block was amplified by PCR and added upstream of the transmembrane and intracellular domains of human CD28 (aa 153-220) followed by the intracellular domain of human CD3ζ (aa 52-164). The CD8 leader sequence and HA-tag were inserted upstream of the CD19 scFv to allow for labeling and detection of CAR-expressing cells (**Supplemental Figure S1a**). To make the lentiviral vector, this sequence was inserted downstream of the human ubiquitin-C promoter in the lentiviral plasmid pFUW using Gibson assembly, as previously described (Dai, Xiao, Bryson, Fang, & Wang, 2012).

To make the CD3ζ-only CAR (Z), PCR was used to amplify the sequence from the N-terminal of the CAR construct through the scFv region, as well as the CD3ζ intracellular domain. The codon optimized CD8 transmembrane domain sequence was synthesized by IDT-DNA and inserted between the scFv and CD3ζ PCR products using PCR. The CAR gene fragment was then reinserted into the lentiviral plasmid using Gibson assembly.

Lentiviral vectors were prepared by transient transfection of 293T cells using a standard calcium phosphate precipitation protocol, as described previously (Dai et al., 2012). The viral supernatants were harvested 48 hours post-transfection and filtered through a 0.45 μm filter (Corning, Corning, NY). For transduction, Jurkat T cells were mixed with fresh viral supernatant and centrifuged for 90 minutes at 1050xg at room temperature. A stable CD19-expressing K562 line was generated in a similar way by transducing parental K562 cells with a lentiviral vector encoding the cDNA of human CD19, as described previously (Siriwon et al., 2018).

To get populations of cells that express the transduced protein at different levels, CAR-expressing Jurkat T cells and CD19-expressing K562 cells were sorted into high, medium, and low populations (referred to as Z^Hi^, Z^Med^, Z^Low^, 28Z^Hi^, 28Z^Med^, 28Z^Low^, 19^Hi^, 19^Med^, and 19^Low^, respectively). To do this, the cells were stained with fluorophore-conjugated antibodies. T cells were first stained with anti-HA antibody for 30 minutes at 4°C, followed by three washes with PBS. The cells were then stained with a secondary alexa-647 conjugated anti-rabbit antibody for 15 minutes at 4°C, followed by three more washed with PBS. CD19 cells were stained with PE conjugated anti-CD19 followed by three washed with PBS. All stained cells were then sorted into the three groups using the BD SORP FACS Aria I cell sorter at the USC stem cell flow cytometry core (**Supplemental Figure S1b**).

### T cell stimulation and ppERK analysis

CAR-expressing Jurkat T cells were stimulated by either HA antibody or CD19-expressing cells. For antibody stimulation, 0.1×10^6^ CAR-expressing cells were incubated with various amounts of anti-HA antibody in 200 µl in 96 well plates in a 37°C water bath. For cellular stimulation, 0.1×10^6^ CAR-expressing Jurkat T cells were combined with various concentrations of CD19-expressing K562 cells in 200 µl in 96 well plates. After the cells were mixed, they were centrifuged at 1000xg for 10 seconds before moving directly into a 37°C water bath. Doubly phosphorylated ERK was measured as described previously (Altan-Bonnet & Germain, 2005). Briefly, to fix the intracellular stimulation reactions after a given amount of time, cells were moved to an ice bath, and ice cold 16% paraformaldehyde solution was added to a final concentration of 4% for 20 minutes. The cells were then centrifuged and resuspended in 100% ice-cold methanol. The cells were incubated at -20°C for at least 30 minutes, followed by 3 washes in 200 µl FACS staining buffer (5% FBS in PBS). Cells were then stained with fluorophore conjugated phospho-ERK antibody for 30 minutes at 4°C in the dark, followed by 3 washes with 200 µl PBS. Fluorescence signal was analyzed using the Miltenyi Biotec flow cytometer and all FACS data were analyzed using FlowJo software. Small Jurkat cells were distinguishable from the large K562 target cells based on their low auto-fluorescence in the forward and side scatter channels.

Upon T cell activation, ERK exhibits a digital (on/off) response. Typically, when ERK is measured as a readout of T cell activation, the percent of ERK positive cells in a population is measured. The response time of the population can then be calculated based on the fit of the phosphorylation time course to a standard sigmoidal curve. In this work, we assume the ERK response time is equal to the time it takes to reach half of the maximal level of phosphorylation (EC50). This depends on both the amount of antigen and the amount of CAR expression (**Supplemental Figure S1c**). In our model, we assume that the deterministic differential equations are representative of the average response of the T cell population. Therefore, we directly compare the ERK response time in the model, which represents the response time of an average cell in the population, to the half maximal cellular population response time. This comparison has previously been shown to relate well (Altan-Bonnet & Germain, 2005).

### Experimental data curve fitting

Graphpad Prism was used to fit the phosphorylated ERK response time from our experimental data to a standard sigmoidal curve, using the non-linear regression curve fit function.

## 3 RESULTS

### 3.1 Model of CAR ERK activation

We constructed a computational mechanistic model that describes the early CAR signaling events leading to T cell activation. We specifically predict how the CAR mediates ERK activation through the MAPK pathway. Studies have indicated that, while the activation of CARs and TCRs have different signal initiating components, the signaling events initiated downstream are not significantly different (Harris et al., 2018). Therefore, to construct our model, we combined four mechanistic signaling modules: (I) CAR-specific phosphorylation based on our previously published models, (II) phosphatase activity, (III) a LAT signalosome, and (IV) a MAPK pathway that leads to ERK activation (**Figure 1**). To characterize the model, we first explored signaling primarily through the CD3ζ CAR stimulatory domain. This allowed us to compare our model to previously developed models in the literature, which largely simplify the TCR to account only for the CD3ζ domain.

Module I focuses on LCK autoregulation and its phosphorylation of the CAR intracellular signaling domains and ZAP-70. We first adapted our model of LCK autoregulation and inhibitory phosphorylation by the kinase CSK to reduce the computational complexity (Rohrs, Wang, et al., 2016). The second model in this module was adapted from our previous work to quantify the kinetics of CAR intracellular domain phosphorylation by LCK (Rohrs et al., 2018). Our published computational investigation, paired with novel *in vitro* phospho-proteomic mass spectrometry, specifically identified the phosphorylation rates of individual tyrosine sites. This work also revealed that the addition of CD28 increases the rate of CD3ζ ITAM phosphorylation, which will be used later in the present study to explore how CD28 affects downstream signaling in the T cell activation system. Once CD3ζ ITAMs are doubly phosphorylated, ZAP-70 is able to bind. ZAP-70 can then be phosphorylated by LCK at several sites. This phosphorylation has a variety of effects: holding ZAP-70 in an open conformation, increasing ZAP-70 catalytic activity, and allowing ZAP-70 to dissociate from CD3ζ (Katz et al., 2017; Sjölin-goodfellow et al., 2015).

In module II, we modeled the activity of phosphatases known to play a role in T cell activation. This module influences both modules I and III. We included two main phosphatases that act throughout the whole model of T cell activation: CD45 and SHP1. CD45 is constitutively active in T cells and prevents unstimulated T cell activation. SHP1 activity is induced upon phosphorylation of the TCR. To explore its effects on CAR activation, we included a mechanism of negative feedback through phosphatase SHP1 recruitment, first modeled by Altan-Bonnet and Germain (Altan-Bonnet & Germain, 2005). SHP1 is recruited to singly phosphorylated CD3ζ ITAMs (from module I), where it can be activated by LCK. SHP1 can then dephosphorylate various proteins in the signaling cascades in modules I and III.

Module III, the LAT signalosome, links the output of module I (activated ZAP-70) to the input of module IV (active SOS and PLCg). Module III begins with free and activated ZAP-70 from module I, which is able to phosphorylate LAT. Phosphorylated LAT can bind to adaptor molecules, GADS and Grb2, which in turn bind to other downstream signaling proteins, such as SLP76, and the inputs to module IV, SOS and PLCg. Phosphorylated CD28 can also bind and recruit several of the proteins in the LAT signalosome.

Module IV focuses on MAPK pathway activation. To initiate this pathway, we adapted a model of Ras-GDP to RasGTP conversion by SOS and Ras-GRP from Das *et al*. (Das et al., 2009). Their model details the allosteric regulation of SOS by active Ras, which results in a positive feedback loop that can transform the analog phosphorylation events derived from TCR or CAR activation to a digital ERK response. The RAS-GTP output of this model was used as the input to a MAPK cascade parameterized by Birtwistle *et al.* (Birtwistle et al., 2012), resulting in doubly phosphorylated ERK. Active ERK also feeds back to modules I and II as it can phosphorylate LCK at a protection site, which prevents interactions with the phosphatase SHP1, as first modeled by Altan-Bonnet and Germain (Altan-Bonnet & Germain, 2005).

Together, these modules constitute a mechanistic description of what are thought to be the most important interactions in the binary decision of T cells to activate ERK. Below, we explore the model in detail and make predictions about the mechanisms through which the individual signaling domains on CARs influence the ERK response.

### 3.2 Model parameterization to literature data

We first fit the model parameters to experimental data to obtain a robust mathematical framework to predict T cell activation leading to ERK phosphorylation (**Figure 2)**. We started by refitting our previous model of LCK regulation to reduce the computational complexity and better constrain the parameters, as described in the Methods section. We fit this minimal model of LCK autoregulation and phosphorylation by CSK to five different experimental conditions in the literature (Hui & Vale, 2014). In total, the values of 11 parameters were estimated using 132 experimental data points. **Figure 2a** shows the model fit to experimental data. The median parameter values as well as the standard deviation for 100 best fit parameter sets are shown in **Figure 2b** and are listed in Supplemental Table S1.

The majority of the downstream model parameters come directly from measurements in the literature or from previously published models. However, some of the parameters were not well defined, because they had not been measured experimentally, they had conflicting values after being fit to the specific assumptions of previous models, or they did not account for the two-dimensional nature of the interactions specifically modeled here. This was particularly true of the parameters governing phosphatase activity, which were shown to significantly influence ERK response time in our sensitivity analysis (**Figure 3**). To better constrain these parameters, we fit the model to published measurements obtained using an *in vitro* system of recombinant proteins interacting on a two-dimensional liposomal membrane (Hui et al., 2017). Hui *et al.* used this system, combining twelve proteins involved in T cell activation, to measure nine different protein phosphorylation states in the presence of varying amounts of CD45. We extracted this data (64 data points) and fit seven model parameters, as described in the Methods section. **Figure 2c, top row** shows the model fit to the experimental data collected in the absence of CSK, best fit parameter values and standard deviations are listed in the Supplemental Table S1. To validate this parameterized model, we extracted an additional data set from Hui *et al.* which includes 145 molecules/µm^2^ CSK (64 data points). As the activity of CSK was fit in our minimal LCK phosphorylation model and was not accounted for in the fitting of the Hui *et al.* CD45 dephosphorylation data, this validation provides confidence that combining our minimal LCK phosphorylation model with the larger CD45 dephosphorylation model can accurately reproduce the signaling network (**Figure 2c, bottom row)**. The median parameter values as well as the standard deviation for 100 best fit parameter sets are shown in **Figure 2d**, and are listed in Supplemental Table S1.

**Figure 3:**
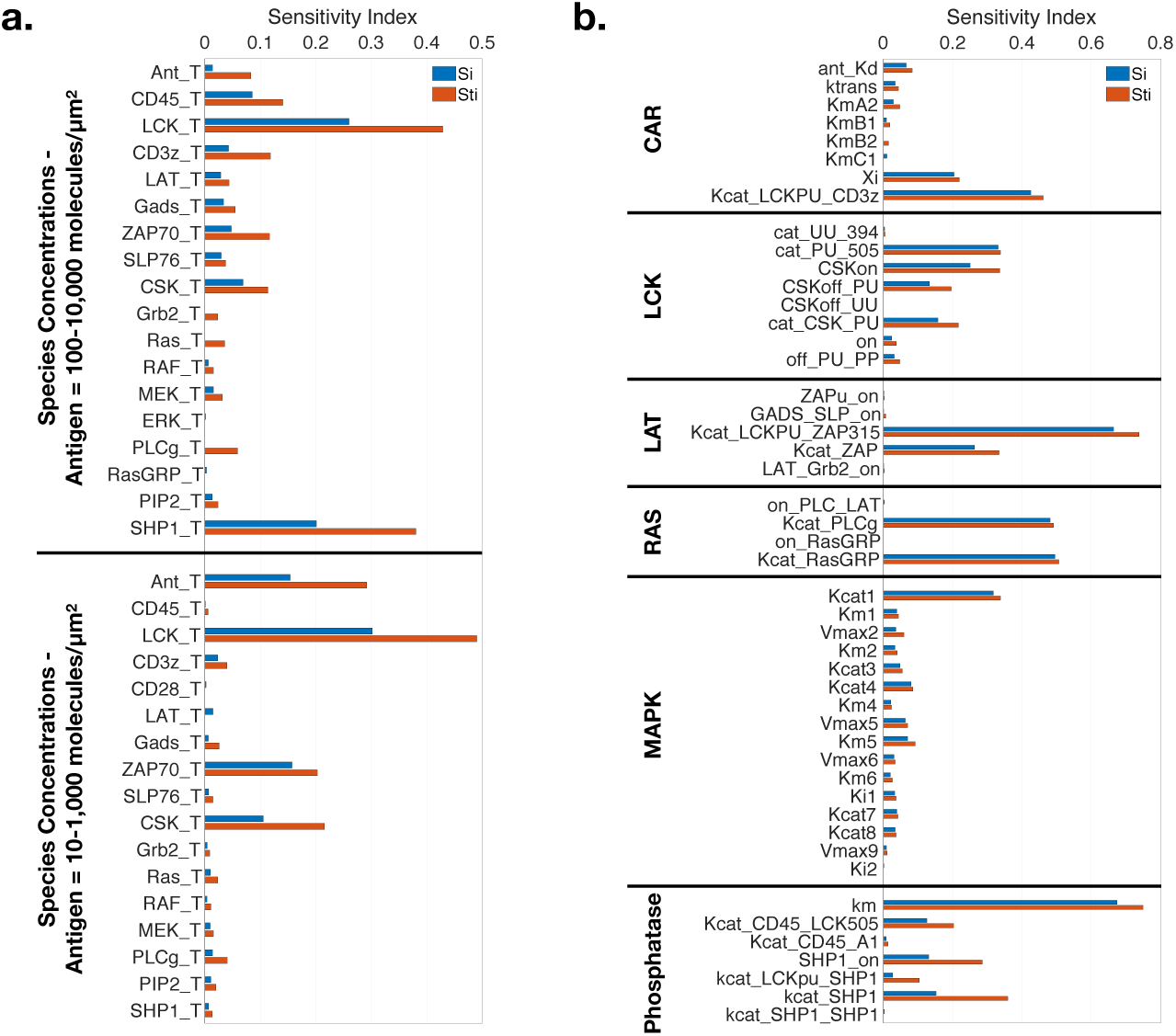
First order (*S*_*i*_) and Total (*S*_*Ti*_) sensitivity indexes of model parameters. a) An eFAST sensitivity analysis was performed on all model initial concentrations for two different nominal ranges of antigen (Ant_T), as indicated on the left. Only the initial conditions whose sensitivity indices are statistically significantly different from that of a dummy variable are shown. The relative sensitivities of species in the model change depending on the amount of antigen in the system. b) Model parameters were separated into nine groups, listed on the left. An eFAST sensitivity analysis was performed on each group with the initial antigen concentration of 100 molecules/µm^2^. Only the parameters whose sensitivity indices are statistically significantly different than that of a dummy variable are shown. For binding interactions with literature defined K_D_ values, only the k_on_ parameter was chosen to vary in the sensitivity analysis.

Overall, the fitted models of LCK autoregulation and phosphatase interactions qualitatively and quantitatively match the experimental data. Additionally, nearly all of the estimated parameter values lie in a tight range. Altogether, we demonstrate that these models recapitulate experiments and can be combined with the other model components to create a predictive framework of CAR-mediated ERK activation. The median values of the estimated parameters were used in model simulations presented below.

### 3.3 Sensitivity analysis reveals network features that control ERK activation

We aimed to first better understand how the model parameters interact with one another and influence the output of doubly phosphorylated ERK response time. To do so, we conducted a sensitivity analysis using the extended Fourier amplitude sensitivity analysis (eFAST) method (Marino et al., 2008). This global sensitivity analysis allows us to identify the parameters that the model output is sensitive to both individually, with the first order sensitivity index (*S*_*i*_), and in combination, with the total sensitivity index (*S*_*Ti*_). This analysis is particularly important for large models, like the one presented here, which incorporate many different mechanisms of feedback and other complex interactions. Parameters with high sensitivity indices strongly influence the model output.

We analyzed the model parameters in seven groups: initial concentrations, CAR parameters, LCK parameters, LAT parameters, RAS parameters, MAPK parameters, and phosphatase parameters. We only list the parameters for which sensitivity indices are statistically significant. Overall, we find that there is at least one parameter in each group that significantly influences ERK response time. Additionally, almost all of the influential parameters have a higher total sensitivity index than first order index. This indicates that, even if all the parameters in a group are not significantly influential on their own, they do all still interact together to affect the output. We next examine the results of the sensitivity analysis in greater detail.

We calculated the sensitivity indices when varying the species’ initial concentrations (**Figure 3a**) for two different conditions, one with a high range of antigen (100-10,000 molecules/µm^2^, top) and one with a moderate range of antigen (1-1,000 molecules/µm^2^, bottom). The relative sensitivity indices of the initial species’ concentrations change between these two experimental conditions. This is particularly interesting when considering the impact of the antigen concentration itself and the concentration of the negative feedback molecule, SHP1. At low antigen concentrations, ERK activation is proportional to the amount of antigen in the system. In this regime LCK, ZAP-70 and CSK emerge as highly influential. This is not entirely surprising, as activation of ZAP-70 is an early bottleneck that must occur before the downstream signaling pathways diverge into more complex branched structures through the many elements of the LAT signalosome. The branches of the LAT signalosome activation converge back onto the MAPK pathway; thus, they are able to help compensate for each other and are less influential overall than the upstream decision makers.

At high antigen concentrations, the sensitivity indices of the antigen concentration are greatly reduced, and the strong influence of SHP1 emerges. We sought to further understand the role of SHP1 and antigen concentration in the model, as the eFAST sensitivity analysis indicated that the interaction between these two species was important. In our model, we assume that the intracellular signaling events downstream of CD3ζ activation are the same for the TCR and CARs. As such, our CAR signaling model incorporates a similar form of SHP1 negative feedback that has been shown to play an important role in TCR signaling. This response has been modeled in TCR signaling previously (Altan-Bonnet & Germain, 2005). We explored this feedback in the model by recording ERK response time for various levels of antigen and SHP1 expression (**Supplemental Figure S2**). As antigen concentration increases for high SHP1 concentrations, as well as in intermediate SHP1 levels (above the red line), the ERK response time first decreases and then increases. When SHP1 concentration is low, this longer ERK response for high CD3ζ is not seen, indicating that it is the feedback of SHP1 that is responsible for this shift in the ERK response time trend. These results reveal that, past a certain threshold antigen concentration, the amount of antigen is not significantly important for controlling the rate of T cell activation. Instead, T cell activation is controlled by the intracellular signaling and negative feedback through SHP1.

The sensitivity indices of these parameters follow similar trends as with the impact of the initial concentrations. **Figure 3b** shows the sensitivity analysis of the other six groups of parameters starting with a moderate concentration of antigen (100 molecules/µm^2^). Some of the most influential parameters in the model are the catalytic rates of LCK, ZAP-70, PLCg, RasGRP, and CD45. Additionally, the value of the Michaelis constant (*km*) is a highly influential parameter since this single parameter plays a role in every Michaelis-Menten reaction in the system. However, there are not enough data to be able to identify both a catalytic rate and Michaelis-Menten constant for all of these reactions, thus leading to our choice to focus our fitting around the highly sensitive catalytic parameters. The calculated sensitivity indices for the network stimulated with a high concentration of antigen (1,000 molecules/µm^2^) shows similar changes as the initial condition sensitivity indices, with SHP1 parameters being more sensitive than the low antigen case and ZAP70 and CSK parameters being slightly less sensitive.

Taken together, the high sensitivity indices of multiple parameters spread throughout the different groups highlights the interconnected nature of the signaling network modeled here, where the final output depends on each step of the pathway to produce a response. Thus, there is not a single category of parameters that solely affects ERK activation. Rather, control of ERK response is distributed across the network.

### 3.4 Model is validated by independent experimental data sets

We next sought to validate the model predictions using experimental data of ERK activation in CAR T cells. In our modeling approach, we assume that the same signaling events that occur downstream of the TCR also occur downstream of the CARs. This assumption has been shown to be true on a macroscale of general phosphorylation events (Harris et al., 2018), and we wanted to further validate it specifically for the negative feedback of SHP1. To do so, we compared model simulations to experimental measurements. First, CAR T cells were made as described in the methods section. We used lentiviral vectors to create stable Jurkat T cell lines expressing HA-tagged anti-CD19 CD28-CD3ζ CARs and sorted them into CAR positive populations (**Supplemental Figure S1**). Using 28z^Med^ Jurkat T cells, we verified that anti-HA antibody is able to bind to the HA-tagged CAR and stimulate ERK phosphorylation. Using this system, we stimulated the cells with various amounts of anti-HA antibody, up to very high concentrations, and measured the percent of doubly phosphorylated ERK over time (**Figure 4a**). We then fit these responses to a 4-parameter sigmoidal curve and estimated the 95% confidence interval of the half maximal ERK response time at each antibody concentration, referred to as the ERK response time (**Figure 4b, black dots and error bars**). For very low concentrations of antibody, the maximal percent of ERK positive cells is also very low, making it difficult to fit a sigmoidal curve. Thus, the confidence intervals for the fitted ERK response times are wide for these low concentrations. However, as we increase the antibody concentration and higher maximal ERK phosphorylation is achieved, the confidence intervals around the fitted ERK response times narrow and we can see a clear trend. As the antibody concentration increases, the ERK response time of the population becomes faster. This trend appears to change at very high antibody concentrations, where the response time begins to slow, presumably, due to the negative feedback from SHP-1, as is seen in endogenous T cell signaling (Altan-Bonnet & Germain, 2005)..

**Figure 4:**
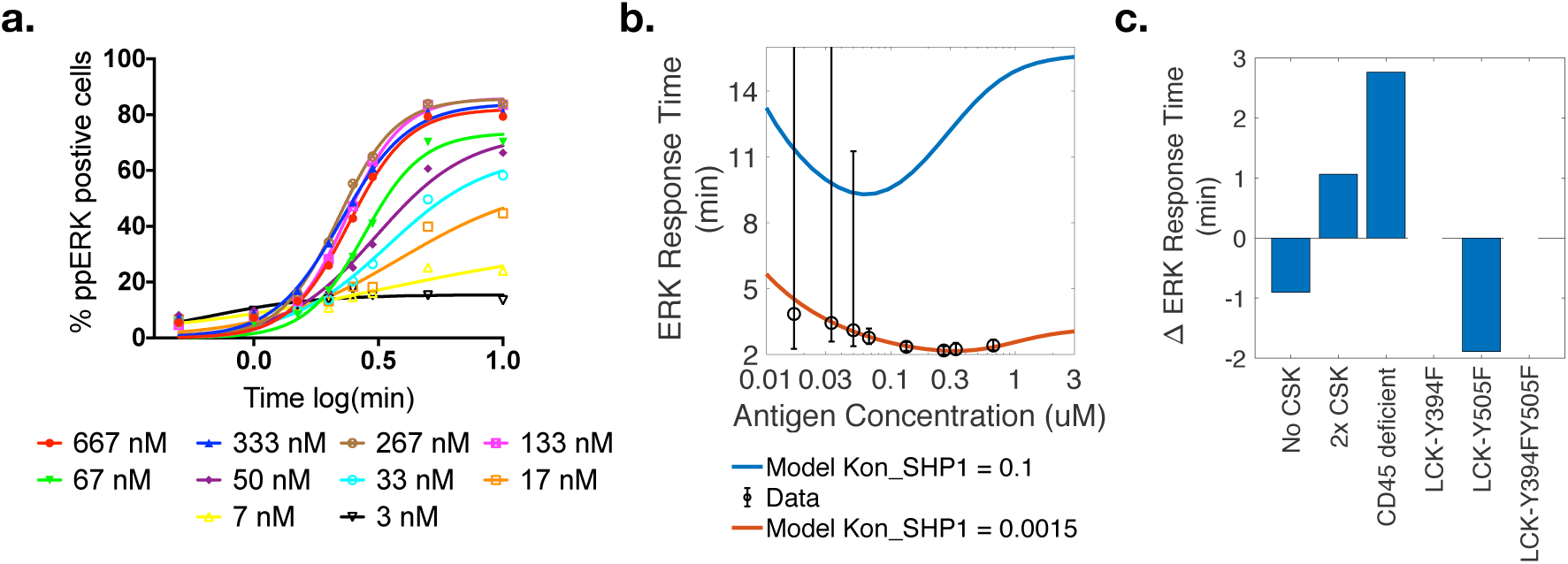
The model can reproduce effects of T cell signaling. a) The percent of ppERK positive 28z^Med^ CAR T cells over time following stimulation with varying amounts of anti-HA antibody. Experimental data (dots) were fit to a sigmoidal curve (lines) to estimate ERK response time (EC50). b) The model simulations (lines) compared the ERK response time of 28z^Med^ CAR T cell activation (dots), calculated from the ppERK response curves in (a). Experimental data is the sigmoidal fit EC50 value from (a) +/-95% confidence interval. Model simulations using the baseline assumption that SHP1 association rate with singly phosphorylated CD3ζ ITAMs is 0.1 µm^2^molecule^-1^min^-1^ (blue line) do not match the data, but changing the SHP1-ITAM association rate to 0.0015 µm^2^molecule^-1^min^-1^ (red line) allows the model to capture the ERK response data well. c) The model can qualitatively match the expected changes in ERK response time due to changes to various intracellular signaling molecules. The change in the ERK response compared to the baseline ERK response model is shown for simulations with varying amounts of CSK as well as alterations to the indicated LCK tyrosine sites to mimic a tyrosine to phenylalanine mutation.

We applied the model to predict the ERK response time for the same antigen concentration levels used in our experiments. Given the mechanistic detail of the model, we could use it to investigate whether SHP1 influences CAR signaling in a similar way as in TCR signaling. Using the baseline model parameters, the model simulations qualitatively agree with the experimental observations, showing faster ERK response time with increasing antigen at low concentrations and slowing ERK response time to a plateau at higher concentrations (**Figure 4b, blue line**). However, the response times given by the model simulations with the baseline parameters are much slower than the experimental data. To address this, we examined to the assumption that all parameters bind with the same association rate (0.1 µm^2^molecule^-1^min^-1^) made during model construction. Based on experimental evidence, we know that SHP1 must be recruited to the T cell signaling area (Lorenz, 2008). As our model does not account for the spatial orientation of molecular diffusion in the cell, we accounted for this step by decreasing the association rate of SHP1 to the singly phosphorylated ITAMs. We tried a range of values for this association rate and found that reducing this rate to 0.0015 µm^2^molecule^-1^min^-1^ allowed the model to match the experimental data. These simulations indicate that SHP1 does indeed play a significant role in CAR signaling. Additionally, we confirm that the model qualitatively and quantitatively matches experimental measurements. We use this reduced association rate in all subsequent model simulations.

We next aimed to further validate the model by determining whether it could qualitatively reproduce known experimental observations obtained following modifications of ERK activation, as published in the literature. To do this, we applied the model to test how different mutations to upstream signaling molecules influence downstream ERK response time (**Figure 4c**). Schoenborn *et al.* modified CSK experimentally to produce a form of the protein that can specifically bind to a small molecule inhibitor (Schoenborn, Tan, Zhang, Shokat, & Weiss, 2011). They showed that inhibiting CSK resulted in faster ERK activation in a population of T cells. When we remove CSK, the model predicts that ERK response time increases by almost one minute, in agreement with the findings from Schoenborn and coworkers. Conversely, when we double the amount of CSK, the model shows that ERK response time slows.

In the same study, Schoenborn *et al.* also showed that CD45 deficient cells have reduced ERK activation upon TCR stimulation. Using the model, we show that removing CD45 greatly slows the ERK response time by roughly 2.75 minutes. These model simulations qualitatively agree with the experimental data.

Similar experiments were done by Philipsen *et al.* to test the ERK response given various LCK tyrosine to phenylalanine mutants expressed in LCK negative Jurkat T cells (Philipsen et al., 2017). They found that LCK-Y394F or LCK-Y394F-Y505F essentially eliminated the ERK positive cell population at three minutes, while LCK-Y505F increased the amount of ERK positive cells. Implementing these two mutations in our model shows that removing LCK-Y394 phosphorylation completely prevents LCK phosphorylation while removing LCK-Y505 phosphorylation speeds up the ERK response time. Since our model does not incorporate stochasticity, we cannot directly measure the percentage of positive cells. However, these trends agree with the experimental findings. Thus, the model is able to capture known effects of signaling modifications in both TCR-and CAR-specific T cell activation.

### 3.5 Model predicts mechanism of CD28-enhanced signaling

The results presented above demonstrate how we have developed a mathematical model to predict ERK activation downstream of CAR signaling. By comparing the model to multiple independent data sets, we present a validated model that can generate reliable, experimentally-based results. This provides confidence that the model can be used to generate new predictions and testable hypotheses.

Therefore, we applied the model to better understand the mechanism of CD28 signaling. In particular, we investigated how the presence of CD28 influences ERK response time, a long-standing question in the field of immunology (Adams, Grierson, Mowat, Harnett, & Garside, 2004; Beyersdorf, Kerkau, & Hünig, 2015). We first quantified how the presence of CD28 affects downstream signaling leading to ERK activation by measuring the ERK response time for Z or 28z CAR T cells. To do this, we expressed the Z CAR in Jurkat T cells, following the same protocol and sorting process used for the 28z Jurkat CAR T cells described earlier. We also expressed CD19 on K562 target cells and sorted them into different expression levels as described in the methods (**Supplemental Figure S1b**). We then stimulated 28z^Med^ and Z^Med^ T cells with different ratios of 19^Med^ target cells and measured the ERK response time (**Figure 5a,b**). Here, we see that the 28z CAR has consistently faster ERK activation for all target cell ratios. To validate that this is a consistent mechanism across a range of CAR and CD19 expression levels, we stimulated 1:1 ratios of high, medium, and low expression CAR and target cells and measured the ERK response time (**Supplemental Figure S1c**). The ERK response time depends on both CAR expression level and CD19 expression level, with high expressing cells displaying faster response times than lower expressing cells. Additionally, 28z CARs had consistently faster ERK response times compared to Z CARs.

**Figure 5:**
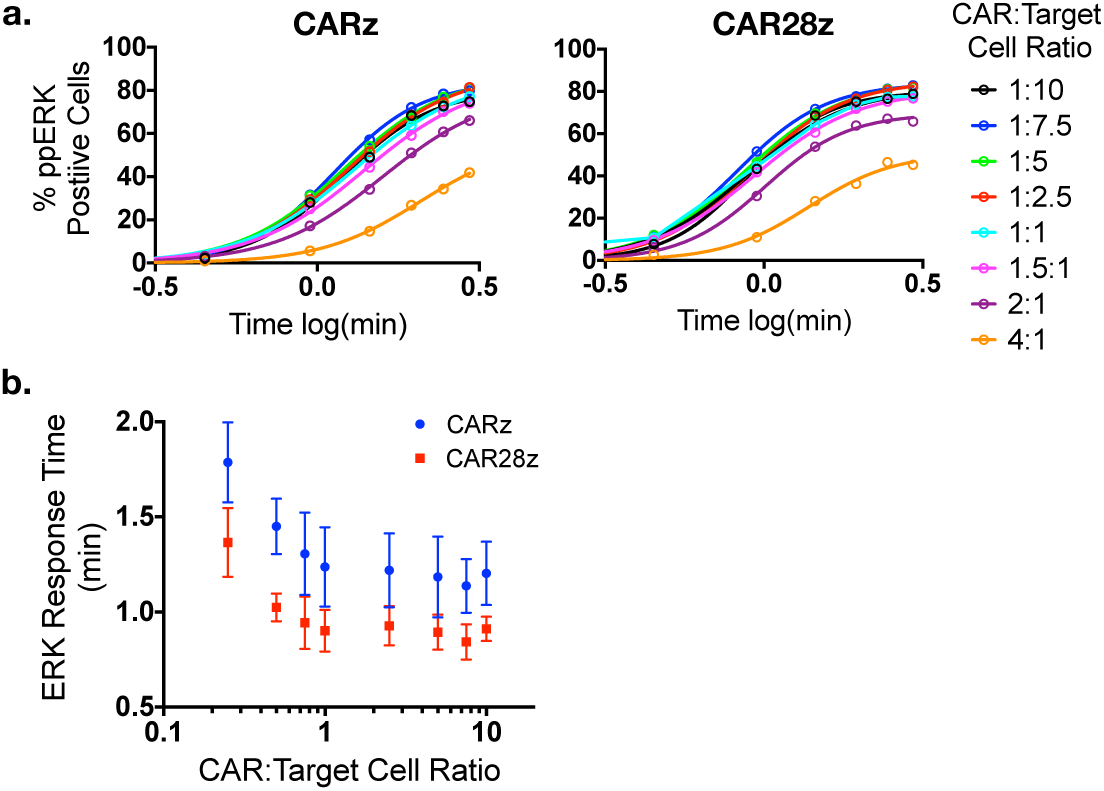
Experimental validation of CAR activated ERK response time. a) Z^Med^ CAR cells and 28z^Med^ CAR cells were mixed with various amounts of 19^Med^ K562 cells, and the ERK response was measured over time. The data (dots) were then fit to a sigmoidal curve (lines), and the ERK response time (EC50) was calculated. b) Experimental ERK response time of Z^Med^ and 28z^Med^ CAR T cell activation (dots). Experimental data are the EC50 value from the sigmoidal curves fit in (A) +/-95% confidence interval.

To understand the mechanisms that lead to faster ERK response time in the presence of CD28, we explored the literature to identify the important mechanisms of CD28 signaling. CD28 has been shown to bind to several different proteins that are also involved in the LAT signalosome. Specifically, phosphorylated tyrosine sites and proline-rich regions on CD28 can bind to the adaptor proteins Grb2 and GADS (Higo et al., 2014). These proteins can recruit and bind to other proteins that lead to ERK activation. Additionally, our previous work to quantify CAR phosphorylation kinetics showed that the presence of CD28, without any downstream binding proteins, increases the rate of CD3ζ phosphorylation (Rohrs et al., 2018). It is possible that enhancing CD3ζ phosphorylation is another mechanism by which CD28 could influence ERK activation.

To understand the relative importance of each of these three mechanisms (Grb2 binding to CD28, GADS binding to CD28, and CD28-mediated enhancement of LCK activity) on ERK activation in CAR T cells, we used an ensemble modeling approach (Mesecke, Urlaub, Busch, Eils, & Watzl, 2014). The three individual mechanisms implemented in the model are shown in **Figure 6**. (1) Grb2 is able to recruit SOS to the signaling area, which can activate Ras and the MAPK pathway directly (**Figure 6a**) (Schneider, Cai, Prasad, Shoelson, & Rudd, 1995). (2) GADS is able to recruit SLP76, thus increasing the amount of this adaptor protein in the signaling region (**Figure 6b**) (Sela et al., 2011; Wonerow & Watson, 2001). For Grb2 and GADS binding, we assume that these adaptor proteins will bind and signal in the same way that they do on the LAT signalosome. (3) The third mechanism uses the kinetic rates calculated in our previous model of phosphorylation of the individual CD3ζ ITAM sites in the presence of CD28. In this mechanism, the increased phosphorylation rate of CD3ζ allows for faster recruitment of ZAP-70 and therefore faster activation of the LAT signalosome and the MAPK pathway (**Figure 6c**) (Rohrs et al., 2018).

**Figure 6:**
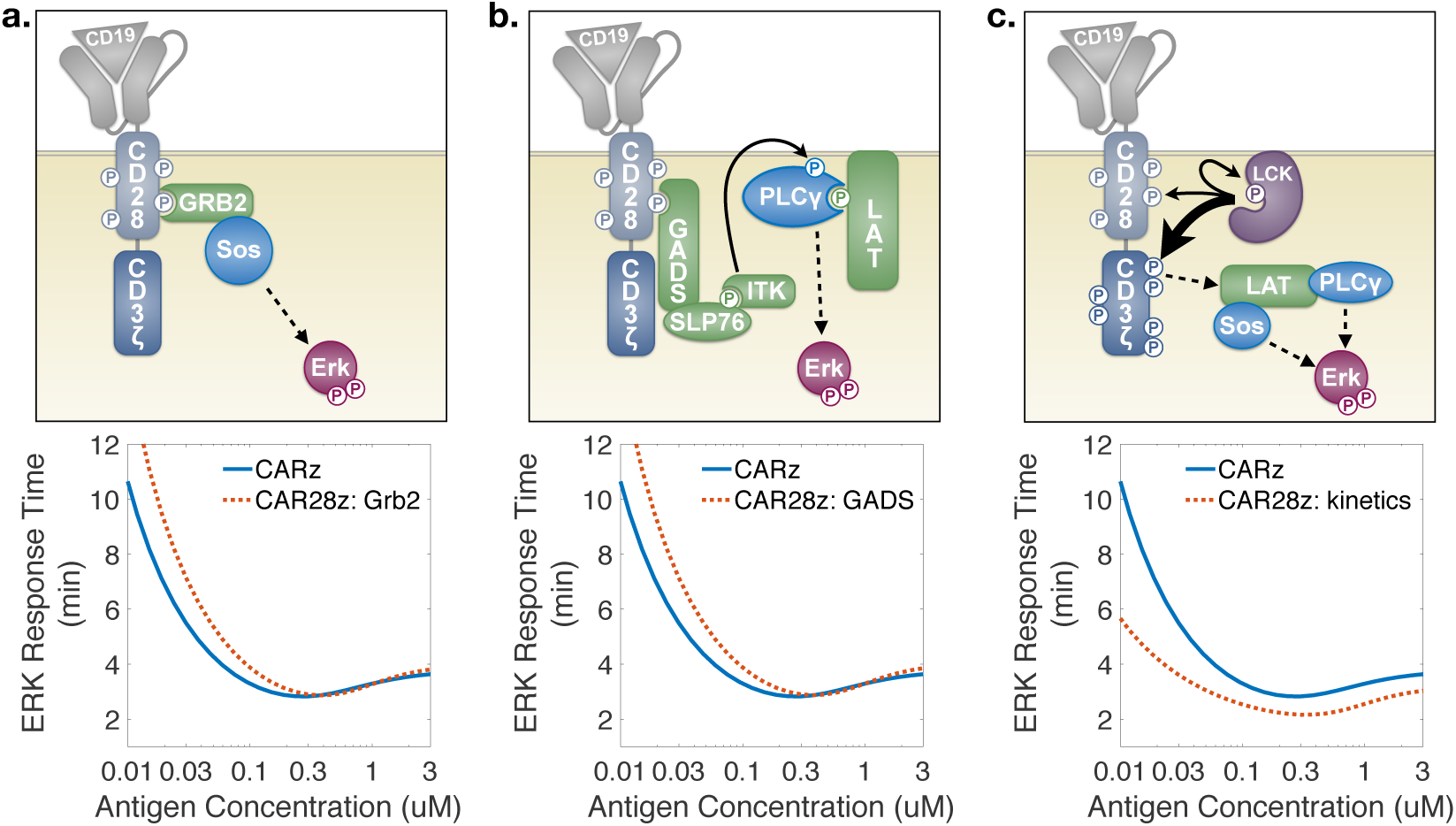
Ensemble models of CD28 ERK activation mechanisms. a) *Top*, CD28 can bind to Grb2, which can bind to SOS and activate the MAPK pathway and ERK. *Bottom*, ERK response time as a function of CD3ζ concentration for the Z (blue) or 28z (red) CAR in which the only effect of CD28 activation is its binding to Grb2. b) *Top*, CD28 can bind to GADS, which can potentially bind to SLP76. Tec family kinases recruited by SLP76 can then activate PLCg on the LAT signalosome, which activates MAPK pathway and ERK. *Bottom*, ERK response time as a function of CD3ζ concentration for the Z (blue) or 28z (red) CAR in which the only effect of CD28 activation is its binding to GADS.

We simulated the model with each CD28 mechanism individually and in various combinations. The bottom panels of **Figure 6** show the predicted ERK response time as a function of antigen concentration for each mechanism individually with CD28 present, compared to the simulated case where the CAR only expresses the CD3ζ domain. Both Grb2 and GADS binding showed similar effects: slightly slowing ERK response time at low antigen concentrations, with only minor effects at high antigen concentrations. In contrast, the effect of the increased rate of CD3ζ phosphorylation was significantly different, showing a nearly constant decrease in response time in the presence of CD28 over all antigen concentrations. Simulating Grb2 and GADS binding together did not appear qualitatively different than simulations with each one individually (**Supplemental Figure S3a**). Adding either or both binding mechanisms to the mechanism of increased LCK kinetics was not significantly different from the increased kinetics mechanism alone (**Supplemental Figure S3b-d**).

Comparing these results to the experimental data in **Figure 5b**, we can see that the model in which CD28 influences the kinetics of LCK phosphorylation of CD3ζ qualitatively matches the experimental data. The difference in the absolute quantification of ERK response times between the target cell stimulated data and the model is likely due to the fact that the model was fit to experimental data of CAR stimulation through anti-HA antibodies, which are expected to be less efficient at inducing the strong crosslinking that aids in immunological synapse formation and more efficient T cell activation induced by cell-cell interactions. However, the model qualitatively predicts the qualitative effects of CAR stimulation, without any additional parameter fitting or tuning.

Therefore, based on these detailed model simulations, we hypothesize that CD28 primarily influences ERK activation through recruitment of LCK, which increases the kinetics of CD3ζ activation, and not through specific binding events of the CD28 protein itself. This hypothesis is validated by the fact that our modeling results closely match the experimental measurements.

## 4 DISCUSSION

In this study, we developed a computational mechanistic model of the signaling events that lead to activation of CAR-engineered T cells via MAPK signaling. To our knowledge, this is the first model to combine this level of detail of the T cell activation signaling and co-stimulatory pathways. The model incorporates 23 different proteins in the signaling pathway that leads from CAR-antigen binding to ERK activation. Experiments quantifying ERK activation in CAR-bearing Jurkat T cells were used for model parameterization and validation. The validated model was used to explore CAR signaling and how the CD28 co-stimulatory domain influences ERK activation. We used an ensemble modeling approach to generate novel hypotheses for the way in which three different CD28 signaling mechanisms influence ERK activation kinetics (Mesecke, Urlaub, Busch, Eils, & Watzl, 2011). Specifically, we confirmed the importance of SHP1 negative feedback in CAR signaling and generated new hypotheses regarding the role of CD28. We show that CD28 primarily affects downstream signaling through recruiting LCK to modify the phosphorylation rate of CD3ζ and that the binding properties of CD28 alone may actually retard T cell activation at low antigen concentrations. These hypotheses can be used in the future to design new experiments to improve our understanding of these engineered proteins.

The model was first parameterized based on estimated values from experimental measurements and previous models in the literature (Altan-Bonnet & Germain, 2005; Birtwistle et al., 2012; Das et al., 2009; Higo et al., 2014; Houtman et al., 2004; Rohrs, Wang, et al., 2016). We then performed a global sensitivity analysis to determine which parameters most strongly influence the ERK response time. From this analysis, we found that the upstream parameters controlling catalytic rates of LCK, ZAP-70, and the phosphatase CD45 were particularly important in influencing the ERK response time. These parameters were not well defined in the literature; however, we highlight their importance in this signaling pathway. Our results provide motivation to better determine the values of those parameters experimentally in the future.

Once the model was fully parameterized, we ensured that it could reproduce experimental CAR-specific T cell activation data, as well as observations for TCR-stimulated ERK activation in the literature. Tuning the SHP1 association rate with singly phosphorylated ITAMs allowed for our model to fit ERK activation data for anti-HA antibody stimulated 28z CAR T cells. The model was also able to capture the effects of various signaling modifications on T cell ERK activation, indicating that the model is robust. Finally, model predictions qualitatively comparing Z and 28z CAR T cell activation were validated by additional experiments.

Given the mechanistic detail of the model, we can distinguish the possible ways that the CD28 co-stimulatory domain affects ERK response time in engineered T cells. CD28 is known to bind to several different adaptor proteins that can recruit activators of Ras and the MAPK pathway (Higo et al., 2014; Tian et al., 2015). Therefore, we investigated how each of these mechanisms may influence the ERK response time. We also tested a finding from our work that CD28 increases the phosphorylation rate of CD3ζ (Rohrs et al., 2018), which in turn could lead to more rapid LAT signalosome formation and ERK activation (Holdorf, Lee, Burack, Allen, & Shaw, 2002). We explored each of these mechanisms alone and in various combinations to develop model-driven hypotheses about how each one would affect the ERK response time. We compared these predictions to experimental data of ERK response time differences between Z and28z CAR T cells stimulated with a 1:1 ratio of CD19-expressing target cells. These experiments qualitatively match the model predictions that the main role of CD28 is to increase CD3ζ phosphorylation kinetics.

The insights from the model increase our understanding of how CD28 is functioning in T cells. The model also generates new hypotheses that can be tested experimentally. Specifically, in the model, the mechanism through which CD28 is able to increase CD3ζ phosphorylation kinetics is not clear. One possibility is that the CD28 domain alters the structure of the CAR on the inner membrane of the T cell to make it more accessible to rapid phosphorylation. Alternatively, CD28 has binding sites for LCK that could be increasing the local concentration of this CD3ζ activating kinase, thus allowing for more rapid phosphorylation. It would be interesting to further test these hypotheses experimentally to more specifically isolate the structural features of CD28 that improve CAR activation. The model also predicts that CD28 binding to GADS and Grb2 may retard T cell ERK activation at low antigen levels. More experimental work is needed to understand the extent of this retardation and how it can be harnessed or modified to improve CAR T cell activation. This iterative approach between hypothesis generation and experimental testing can be used to make more optimal next generation CARs.

We do acknowledge some limitations in the current model. First, the model does not indicate how CD28 influences other downstream T cell activation pathways. In the literature, CD28 has been shown to bind to PI3K, which activates the Akt pathway (Acuto & Michel, 2003; Rudd, Taylor, & Schneider, 2009). Additionally, CD28 co-stimulation with the TCR can increase the amount of active Vav in the T cell (Helou, Petrashen, & Salomon, 2015; Muscolini et al., 2015). These mechanisms are not specifically included in the model, but it is possible that these pathways may cross talk with the MAPK pathway and further influence ERK activation (Costello et al., 1999; Dent, 2014). Additionally, this work does not explore the differences between CD28 signaling when incorporated on the CAR compared to signaling through the traditional separate CD28 molecule, which could have additional implications for dual target CAR therapies (Morello, Sadelain, & Adusumilli, 2016). As new data emerges, the model can be updated to include these alternative mechanisms to help improve our understanding of how CD28 co-stimulatory signaling can be optimized in CAR T cells.

## 5 CONLCUSION

Altogether, the mechanistic model of CAR-mediated T cell signaling we have constructed is able to reproduce known effects of CAR activation of the ERK/MAPK pathway and shed new light on the mechanisms of CAR co-stimulatory signaling through CD28. The model has provided a specific mechanism for the modification of ERK response time by CD28, which matches experimental data. Additionally, the model provides new hypotheses that can be tested experimentally to better understand how to modulate the effects of CD28 signaling in CAR therapies. Thus, the model provides a framework that can be used to better understand and optimize CAR-engineered T cell development.

## ACKNOWLEDGEMENTS

This work was supported by the National Cancer Institute of the National Institutes of Health under Award Numbers F31CA200242 (to J.A.R.), R01EB017206, R01CA170820, and P01CA132681 (to P.W.)

## SUPPORTING INFORMATION

**File S1.** Model parameter values

**File S2.** Model initial concentrations

**File S3.** CD28CD3z BioNetGen model

## AUTHOR CONTRIBUTIONS

J.A.R, S.D.F. and P.W. conceived of the presented idea. J.A.R. planned and carried out computational model development and simulations. J.A.R. planned experiments. J.A.R. and E.S. carried out experiments. S.D.F. and P.W. supervised the project. All authors discussed the results and contributed to the final manuscript.

## REFERENCES

Acuto, O., & Michel, F. (2003). CD28-mediated co-stimulation: a quantitative support for TCR signalling. Nature Reviews. Immunology, 3(12), 939–951. https://doi.org/10.1038/nri1248

Adams, C. L., Grierson, A. M., Mowat, A. M., Harnett, M. M., & Garside, P. (2004). Differences in the kinetics, amplitude, and localization of ERK activation in anergy and priming revealed at the level of individual primary T cells by laser scanning cytometry. Journal of Immunology, 173, 1579–1586. https://doi.org/10.4049/jimmunol.173.3.1579

Altan-Bonnet, G., & Germain, R. N. (2005). Modeling T cell antigen discrimination based on feedback control of digital ERK responses. PLoS Biology, 3(11), e356. https://doi.org/10.1371/journal.pbio.0030356

Arulraj, T., & Barik, D. (2018). Mathematical modeling identifies Lck as a potential mediator for PD-1 induced inhibition of early TCR signaling. PloS One, 13(10), e0206232. https://doi.org/10.1371/journal.pone.0206232

Barua, D., Faeder, J. R., & Haugh, J. M. (2007). Structure-based kinetic models of modular signaling protein function: Focus on Shp2. Biophysical Journal, 92(7), 2290–2300. https://doi.org/10.1529/biophysj.106.093484

Beyersdorf, N., Kerkau, T., & Hünig, T. (2015). CD28 co-stimulation in T-cell homeostasis: a recent perspective. ImmunoTargets and Therapy, 4, 111–122. https://doi.org/10.2147/ITT.S61647

Birtwistle, M. R., Rauch, J., Kiyatkin, A., Aksamitiene, E., Dobrzyński, M., Hoek, J. B., … Kholodenko, B. N. (2012). Emergence of bimodal cell population responses from the interplay between analog single-cell signaling and protein expression noise. BMC Systems Biology, 6, 109. https://doi.org/10.1186/1752-0509-6-109

Bogin, Y., Ainey, C., Beach, D., & Yablonski, D. (2007). SLP-76 mediates and maintains activation of the Tec family kinase ITK via the T cell antigen receptor-induced association between SLP-76 and ITK. Proceedings of the National Academy of Sciences, 104(16), 6638–6643. https://doi.org/10.1073/pnas.0609771104

Braiman, A., Barda-Saad, M., Sommers, C. L., & Samelson, L. E. (2006). Recruitment and activation of PLCgamma1 in T cells: a new insight into old domains. The EMBO Journal, 25(4), 774–784. https://doi.org/10.1038/sj.emboj.7600978

Brännmark, C., Palmér, R., Glad, S. T., Cedersund, G., & Strålfors, P. (2010). Mass and information feedbacks through receptor endocytosis govern insulin signaling as revealed using a parameter-free modeling framework. Journal of Biological Chemistry, 285(26), 20171–20179. https://doi.org/10.1074/jbc.M110.106849

Bretscher, P. A. (1999). A two-step, two-signal model for the primary activation of precursor helper T cells. Proceedings of the National Academy of Sciences of the United States of America, 96(1), 185–190. https://doi.org/10.1073/pnas.96.1.185

Brownlie, R. J., & Zamoyska, R. (2013). T cell receptor signalling networks: branched, diversified and bounded. Nature Reviews. Immunology, 13(4), 257–269. https://doi.org/10.1038/nri3403

Bu, J., Shaw, A. S., & Chanti, A. C. (1995). Analysis of then interaction of Zap-70 and Syk protein-tyrosine kinases with the T cell antigen receptor by plasmon resonance. PNAS, 92(May), 5106–5110.

Coombs, D., & Goldstein, B. (2005). T cell activation: Kinetic proofreading, serial engagement and cell adhesion. Journal of Computational and Applied Mathematics, 184(1), 121–139. https://doi.org/10.1016/j.cam.2004.07.035

Costello, P. S., Walters, a E., Mee, P. J., Turner, M., Reynolds, L. F., Prisco, a, … Tybulewicz, V. L. (1999). The Rho-family GTP exchange factor Vav is a critical transducer of T cell receptor signals to the calcium, ERK, and NF-kappaB pathways. Proceedings of the National Academy of Sciences of the United States of America, 96(6), 3035–3040. https://doi.org/10.1073/pnas.96.6.3035

Dai, B., Xiao, L., Bryson, P. D., Fang, J., & Wang, P. (2012). PD-1/PD-L1 blockade can enhance HIV-1 gag-specific T cell immunity elicited by dendritic cell-directed lentiviral vaccines. Molecular Therapy, 20(9), 1800–1809. https://doi.org/10.1038/mt.2012.98

Das, J., Ho, M., Zikherman, J., Govern, C., Yang, M., Weiss, A., … Roose, J. P. (2009). Digital signaling and hysteresis characterize ras activation in lymphoid cells. Cell, 136(2), 337–351. https://doi.org/10.1016/j.cell.2008.11.051

Davis, S. J., & van der Merwe, P. A. (2006). The kinetic-segregation model: TCR triggering and beyond. Nature Immunology, 7(8), 803–809. https://doi.org/10.1038/ni1369

Dent, P. (2014). Crosstalk between ERK, AKT, and cell survival. Cancer Biology and Therapy, 15(3), 245–246. https://doi.org/10.4161/cbt.27541

Harris, D. T., Hager, M. V, Smith, S. N., Stone, J. D., Kruger, P., Lever, M., … Greenberg, P. D. (2018). Comparison of T Cell Activities Mediated by Human TCRs and CARs That Use the Same Recognition Domains. Journal of Immunology. https://doi.org/10.4049/jimmunol.1700236

Helou, Y. a., Petrashen, A. P., & Salomon, A. R. (2015). Vav1 regulates T cell activation through a feedback mechanism and crosstalk between the T cell receptor and CD28. Journal of Proteome Research, 150604190820001. https://doi.org/10.1021/acs.jproteome.5b00340

Higo, K., Oda, M., Morii, H., Takahashi, J., Harada, Y., Ogawa, S., & Abe, R. (2014). Quantitative analysis by surface plasmon resonance of CD28 interaction with cytoplasmic adaptor molecules Grb2, Gads and p85 PI3K. Immunological Investigations, 1–14. https://doi.org/10.3109/08820139.2013.875039

Holdorf, A. D., Lee, K.-H., Burack, W. R., Allen, P. M., & Shaw, A. S. (2002). Regulation of Lck activity by CD4 and CD28 in the immunological synapse. Nature Immunology, 3(3), 259–264. https://doi.org/10.1038/ni761

Houtman, J. C. D., Higashimoto, Y., Dimasi, N., Cho, S., Yamaguchi, H., Bowden, B., … Samelson, L. E. (2004). Binding specificity of multiprotein signaling complexes is determined by both cooperative interactions and affinity preferences. Biochemistry, 43(14), 4170–4178. https://doi.org/10.1021/bi0357311

Hui, E., Cheung, J., Zhu, J., Su, X., Taylor, M. J., Wallweber, H. A., … Vale, R. D. (2017). T cell costimulatory receptor CD28 is a primary target for PD-1–mediated inhibition. Science, 4(March), eaaf1292. https://doi.org/10.1126/science.aaf1292

Hui, E., & Vale, R. D. (2014). In vitro membrane reconstitution of the T-cell receptor proximal signaling network. Nature Structural & Molecular Biology, 21(2), 133–142. https://doi.org/10.1038/nsmb.2762

Huppa, J. B., & Davis, M. M. (2003). T-cell-antigen recognition and the immunological synapse. Nature Reviews Immunology, 3(12), 973–983. https://doi.org/10.1038/nri1245

Iadevaia, S., Lu, Y., Morales, F. C., Mills, G. B., & Ram, P. T. (2010). Identification of optimal drug combinations targeting cellular networks: integrating phospho-proteomics and computational network analysis. Cancer Research, 70(17), 6704–6714. https://doi.org/10.1158/0008-5472.CAN-10-0460

Katz, Z. B., Novotná, L., Blount, A., & Lillemeier, B. F. (2017). A cycle of Zap70 kinase activation and release from the TCR amplifies and disperses antigenic stimuli. Nature Immunology, 18(1). https://doi.org/10.1038/ni.3631

Leupin, O., Zaru, R., Laroche, T., Müller, S., & Valitutti, S. (2000). Exclusion of CD45 from the T-cell receptor signaling area in antigen-stimulated T lymphocytes. Current Biology, 10(5), 277–280. https://doi.org/10.1016/S0960-9822(00)00362-6

Lipniacki, T., Hat, B., Faeder, J. R., & Hlavacek, W. S. (2008). Stochastic effects and bistability in T cell receptor signaling. Journal of Theoretical Biology, 254(1), 110–122. https://doi.org/10.1016/j.jtbi.2008.05.001

Lorenz, U. (2008). Shp-1, Shp-2. Science and Children, 46(2), 26–29. https://doi.org/10.1111/j.1600-065X.2008.00760.x.SHP-1

Marino, S., Hogue, I. B., Ray, C. J., & Kirschner, D. E. (2008). A methodology for performing global uncertainty and sensitivity analysis in systems biology. Journal of Theoretical Biology, 254(1), 178–196. https://doi.org/10.1016/j.jtbi.2008.04.011

McKeithan, T. W. (1995). Kinetic proofreading in T-cell receptor signal transduction. Proceedings of the National Academy of Sciences of the United States of America, 92(11), 5042–5046. Retrieved from http://www.pubmedcentral.nih.gov/articlerender.fcgi?artid=41844&tool=pmcentrez&rendertype=abstract

Mesecke, S., Urlaub, D., Busch, H., Eils, R., & Watzl, C. (2011). Integration of activating and inhibitory receptor signaling by regulated phosphorylation of Vav1 in immune cells. Science Signaling, 4(175), 1–10. https://doi.org/10.1126/scisignal.2001325

Mesecke, S., Urlaub, D., Busch, H., Eils, R., & Watzl, C. (2014). Integration of Activating and Inhibitory Receptor Signaling by Regulated Phosphorylation of Vav1 in Immune Cells. Science Signaling, 4(175). https://doi.org/10.1126/scisignal.2001325

Milone, M. C., Fish, J. D., Carpenito, C., Carroll, R. G., Binder, G. K., Teachey, D., … June, C. H. (2009). Chimeric receptors containing CD137 signal transduction domains mediate enhanced survival of T cells and increased antileukemic efficacy in vivo. Molecular Therapy: The Journal of the American Society of Gene Therapy, 17(8), 1453–1464. https://doi.org/10.1038/mt.2009.83

Morello, A., Sadelain, M., & Adusumilli, P. S. (2016). Mesothelin-targeted CARs: Driving T cells to solid Tumors. Cancer Discovery, 6(2), 133–146. https://doi.org/10.1158/2159-8290.CD-15-0583

Morgan, R. A., Yang, J. C., Kitano, M., Dudley, M. E., Laurencot, C. M., & Rosenberg, S. A. (2010). Case report of a serious adverse event following the administration of t cells transduced with a chimeric antigen receptor recognizing ERBB2. Molecular Therapy, 18(4), 843–851. https://doi.org/10.1038/mt.2010.24

Mukherjee, M., Mace, E. M., Carisey, A. F., Ahmed, N., & Orange, J. S. (2017). Quantitative Imaging Approaches to Study the CAR Immunological Synapse. Molecular Therapy, 25(8), 1757–1768. https://doi.org/10.1016/j.ymthe.2017.06.003

Mullard, A. (2017). Second anticancer CAR T therapy receives FDA approval. Nature Reviews Drug Discovery, 16(12), 818–818. https://doi.org/10.1038/nrd.2017.249

Muscolini, M., Camperio, C., Porciello, N., Caristi, S., Capuano, C., Viola, A., … Tuosto, L. (2015). Phosphatidylinositol 4-phosphate 5-kinase alpha and Vav1 mutual cooperation in CD28-mediated actin remodeling and signaling functions. J Immunol, 194(3), 1323–1333. https://doi.org/10.4049/jimmunol.1401643

Nag, A., Monine, M. I., Faeder, J. R., & Goldstein, B. (2009). Aggregation of membrane proteins by cytosolic cross-linkers: theory and simulation of the LAT-Grb2-SOS1 system. Biophysical Journal, 96(7), 2604–2623. https://doi.org/10.1016/j.bpj.2009.01.019

Northrup, S. H., & Erickson, H. P. (1992). Kinetics of protein-protein association explained by Brownian dynamics computer simulation. Proceedings of the National Academy of Sciences of the United States of America, 89(April), 3338–3342. https://doi.org/10.1073/pnas.89.8.3338

Philipsen, L., Reddycherla, A. V., Hartig, R., Gumz, J., Kästle, M., Kritikos, A., … Müller, A. J. (2017). De novo phosphorylation and conformational opening of the tyrosine kinase Lck act in concert to initiate T cell receptor signaling. Science Signaling, 10(462). https://doi.org/10.1126/scisignal.aaf4736

Rohrs, J. A., Sulistio, C. D., & Finley, S. D. (2016). Predictive model of throm-bospondin-1 and vascular endothelial growth factor in breast tumor tissue. Npj Systems Biology And Applications, 2, 16030. Retrieved from http://dx.doi.org/10.1038/npjsba.2016.30

Rohrs, J. A., Wang, P., & Finley, S. D. (2016). Predictive Model of Lympho-cyte-Specific Protein Tyrosine Kinase (LCK) Autoregulation. Cellular and Molecular Bioengineering, 9(3), 351–367. https://doi.org/10.1007/s12195-016-0438-7

Rohrs, J. A., Wang, P., & Finley, S. D. (2019). Understanding the Dynamics of T-Cell Activation in Health and Disease Through the Lens of Computational Modeling. JCO Clinical Cancer Informatics, (3), 1–8. https://doi.org/10.1200/cci.18.00057

Rohrs, J. A., Zheng, D., Graham, N. A., Wang, P., & Finley, S. (2018). Computational model of chimeric antigen receptors explains site-specific phosphorylation kinetics. BioRxiv. Retrieved from http://biorxiv.org/content/early/2018/02/08/262527.abstract

Rudd, C. E., Taylor, A., & Schneider, H. (2009). CD28 and CTLA-4 coreceptor expression and signal transduction. Immunological Reviews, 229(1), 12–26. https://doi.org/10.1111/j.1600-065X.2009.00770.x

Sadelain, M., Brentjens, R., & Rivière, I. (2013). The basic principles of chimeric antigen receptor design. Cancer Discovery, 3(4), 388–398. https://doi.org/10.1158/2159-8290.CD-12-0548

Schlosshauer, M., & Baker, D. (2004). Realistic protein-protein association rates from a simple diffusional model neglecting long-range interactions, free energy barriers, and landscape ruggedness. Protein Science: A Publication of the Protein Society, 13(6), 1660–1669. https://doi.org/10.1110/ps.03517304

Schneider, H., Cai, Y.-C, Prasad, K. V. S., Shoelson, S. E., & Rudd, C. E. (1995). T cell antigen CD28 binds to the GRB-2/SOS complex, regulators of p21ras. European Journal of Immunology, 25(4), 1044–1050. https://doi.org/10.1002/eji.1830250428

Schoenborn, J. R., Tan, Y. X., Zhang, C., Shokat, K. M., & Weiss, A. (2011). Feedback circuits monitor and adjust basal Lck-dependent events in T cell receptor signaling. Science Signaling, 4(190), 1–14. https://doi.org/10.1126/scisignal.2001893

Sela, M., Bogin, Y., Beach, D., Oellerich, T., Lehne, J., Smith-Garvin, J. E., … Yablonski, D. (2011). Sequential phosphorylation of SLP-76 at tyrosine 173 is required for activation of T and mast cells. The EMBO Journal, 30(15), 3160–3172. https://doi.org/10.1038/emboj.2011.213

Siriwon, N., Kim, Y. J., Siegler, E., Chen, X., Rohrs, J. A., Liu, Y., & Wang, P. (2018). CAR-T Cells Surface-Engineered with Drug-Encapsulated Nanoparticles Can Ameliorate Intratumoral T-cell Hypofunction. Cancer Immunology Research, 6(7), 812–824. https://doi.org/10.1158/2326-6066.CIR-17-0502

Sjölin-goodfellow, H., Frushicheva, M. P., Ji, Q., Cheng, D. A., Kadlecek, T. A., Cantor, A. J., & Kuriyan, J. (2015). The catalytic activity of the kinase ZAP-70 mediates basal signaling and negative feedback of the T cell receptor pathway. Science Signaling, 8(377), 1–14.

Tian, R., Wang, H., Gish, G. D., Petsalaki, E., Pasculescu, A., Shi, Y., … Pawson, T. (2015). Combinatorial proteomic analysis of intercellular signaling applied to the CD28 T-cell costimulatory receptor. Proc Natl Acad Sci U S A, 112(13), E1594–603. https://doi.org/10.1073/pnas.1503286112

Watanabe, K., Kuramitsu, S., Posey, A. D., & June, C. H. (2018). Expanding the therapeutic window for CAR T cell therapy in solid tumors: The knowns and unknowns of CAR T cell biology. Frontiers in Immunology, 9(OCT), 1–12. https://doi.org/10.3389/fimmu.2018.02486

Wonerow, P., & Watson, S. P. (2001). The transmembrane adapter LAT plays a central role in immune receptor signalling. Oncogene, 20, 6273–6283. https://doi.org/10.1038/sj.onc.1204770

