## Supplemental Figure for "ERK activation in CAR T cells is amplified by CD28-mediated increase in CD3ζ phosphorylation"

### Supplemental Figures:

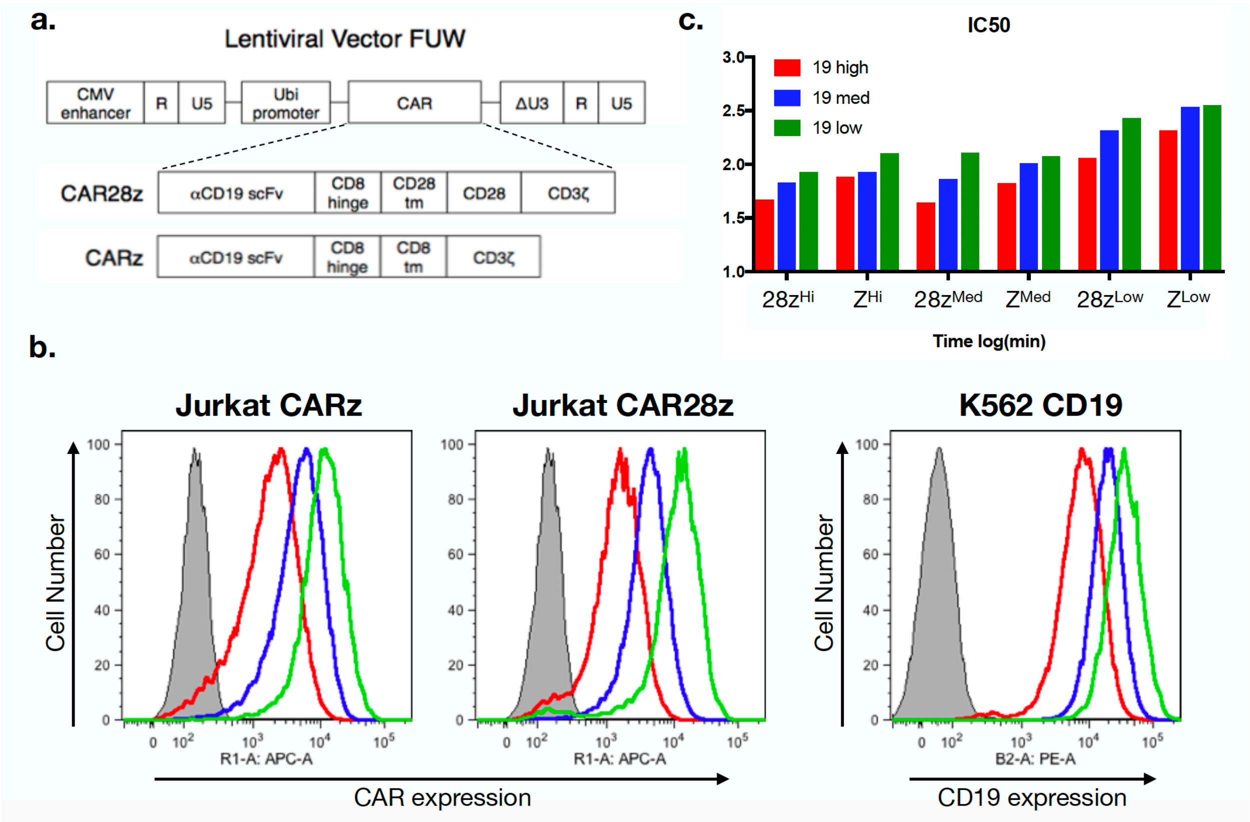


**Supplemental Figure S1: CAR and CD19 expression and T cell activation.**

a) Schematic of the CAR lentiviral vectors. Sequences encoding anti-CD19 CARs were inserted downstream of the Ubi promoter in the FUW lentiviral vector.

b) (Left, middle) CAR expression in Jurkat T cells. CARs were expressed using lentiviral vectors and the resulting populations were sorted into high (green), medium (blue) and low (red) expressing populations. The grey area shows the isotype control staining. (Right) CD19 expression in K562 cells. CD19 was expressed using a lentiviral vector and the resulting population was sorted into high (green), medium (blue) and low (red) expressing populations. The grey area shows the isotype control staining.

c) The ERK response time with varying populations of CD19-expressing cells. High, medium, and low CAR-expressing cell populations were mixed at a 1:1 ratio with high, medium, and low CD19 K562 cell populations, and the percentage of ERK positive cells was measured over time. A sigmoidal curve was fit to the time course data, and the ERK response time (EC50) was calculated.

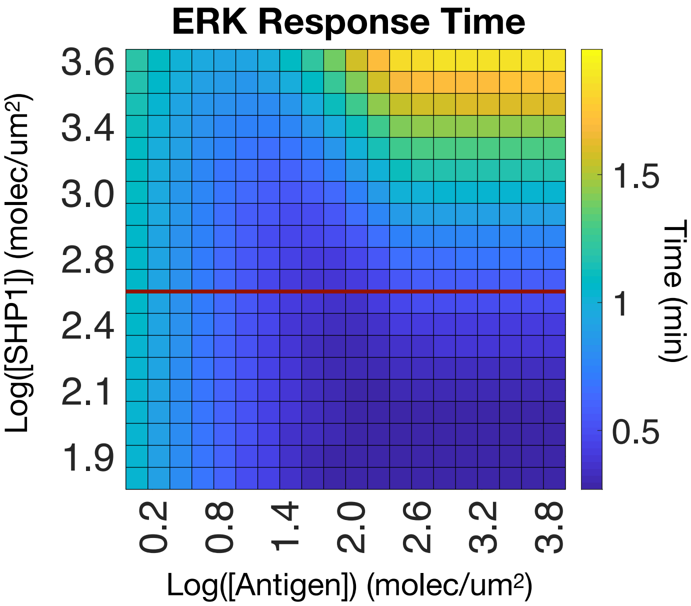


**Supplemental Figure S2: Effect of SHP1 on ppERK CAR T cell activation**. Antigen and SHP1 concentrations were varied in the model and the ERK response time was calculated. The concentration of SHP1 which qualitatively matches the experimental data in Figure 4B is shown in red.

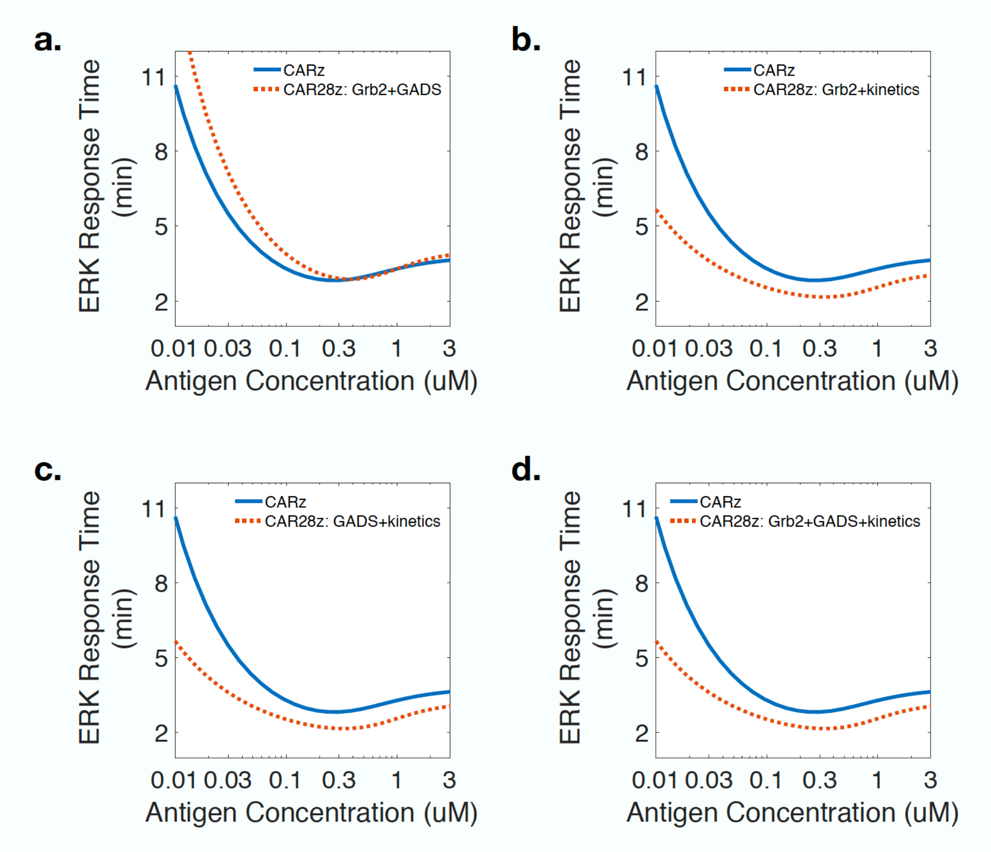


**Supplemental Figure S3: CD28 mechanism ensemble modeling combination simulations**.

a) ERK response time as a function of CD3ζ concentration for the Z (blue) or 28z (red) CAR in which CD28 activation allows for both Grb2 and GADS to bind.

b) ERK response time as a function of CD3ζ concentration for the Z (blue) or 28z (red) CAR in which the only effect of CD28 activation allows for both Grb2 to bind and for the increased rate of CD3ζ phosphorylation.

c) ERK response time as a function of CD3ζ concentration for the Z (blue) or 28z (red) CAR in which the only effect of CD28 activation allows for both GADS to bind and for the increased rate of CD3ζ phosphorylation.

d) ERK response time as a function of CD3ζ concentration for the Z (blue) or 28z (red) CAR in which the only effect of CD28 activation allows for both GADS and Grb2 to bind as well as increasing rate of CD3ζ phosphorylation.
